# Processing of inner bodily signals: evidence and insight from adolescence

**DOI:** 10.1101/2025.08.01.668055

**Authors:** Silvia Canino, Valentina Torchia, Erica Dolce, Irene Ruffo, Teresa Iona, Simona Raimo, Liana Palermo

**Affiliations:** Department of Medical and Surgical Sciences, Magna Graecia University of Catanzaro, Catanzaro, Italy; Department of Health Sciences, Magna Graecia University of Catanzaro, Catanzaro, Italy

## Abstract

Interoception, the sense of inner bodily signals, plays a key role in emotional regulation, cognition and mental health. While its relevance in adulthood has been extensively explored, less is known about how these abilities develop during adolescence, a period characterised by significant physical and psychological changes. This study aimed to investigate three distinct dimensions of interoception — accuracy, sensitivity and awareness — in adolescents and adults to better understand the developmental profile of this sense.

Fifty-four adolescents (aged 12–14) and 50 adults (aged 25–34) completed the Heartbeat Monitoring Task to assess their actual ability to detect heartbeats, their confidence in this ability, and the confidence-accuracy correspondence, and a questionnaire on the tendency to focus on bodily sensations. The study also examined where participants localised bodily sensations during the interoceptive task.

The results revealed no significant differences in interoceptive accuracy between the two groups. Both age groups exhibited similar body localisation patterns, primarily focusing on the chest during heartbeat detection. However, adolescents showed significantly lower metacognitive awareness of their ability to perceive internal bodily sensations, and higher focus on interoceptive sensations, as reflected in their higher confidence ratings and questionnaire scores. No significant correlations emerged among the three interoceptive dimensions in either group, which supports the view that these dimensions represent independent components of interoception. These findings suggest that, while basic interoceptive detection may be established by early adolescence, the capacity to accurately reflect on these internal sensations continues to mature into adulthood. The mismatch observed between adolescents’ heightened bodily focus and their limited metacognitive insight may partly help explain why adolescence is a period of increased vulnerability to mental health difficulties.

## Introduction

Interoception, the sense of the physiological condition of the inner body, is integral to how we experience and interpret bodily signals [1,2]. This sense encompasses sensations and representations of physiological signals, such as the heartbeat, itchiness and air hunger [1]. Research has highlighted the importance of interoception in emotional regulation, cognition and overall well-being [3-8].

On a conscious level, interoception can be operationalized along three main dimensions: (i) interoceptive accuracy (IAcc), which refers to performance on objective tasks like heartbeat detection; (ii) interoceptive sensibility (ISe), which is the self-evaluated tendency to focus on interoceptive signals, measured through questionnaires; and (iii) interoceptive awareness (IAw), which is the metacognitive ability to assess how accurately one perceives internal signals, evaluated through the correspondence between confidence and actual performance [9]. This taxonomy is further supported by evidence suggesting differential contributions of these dimensions on cognition (e.g., [10, 3]) and their different relations with mental health difficulties (e.g., [11, 12]).

Despite the growing recognition of interoception’s role in shaping psychological functioning, as highlighted in a seminal review on interoceptive development by Murphy et al. [13], our understanding of how interoception develops across the lifespan remains limited. In particular, research explicitly investigating interoception during typical adolescence is still scarce (for a similar argument, see also [14]).

Adolescence, however, is a period of life that can be particularly relevant for interoceptive learning, as it is characterised by significant body changes [13, 15]. Adolescence is also marked by significant maturation of neural circuits involved in processing internal bodily signals, influencing how adolescents perceive and respond to internal states such as hunger, fatigue, and emotional arousal [16]. The maturation of interoceptive processes during this period seems to be linked to the development of self-regulation and emotional resilience [17]. Also, disruptions in interoceptive processing during adolescence can contribute to the onset of mental health disorders, such as anxiety and depression [6].

Recent neurophysiological studies have started to explore how interoception manifests in the adolescent brain. For example, Mai et al. [18] showed that heartbeat-evoked potentials (HEPs), a neural marker of interoceptive processing, are associated with an IAcc measure but not with an ISe measure in adolescents, providing objective evidence of the neurocognitive underpinnings of bodily awareness in this age group.

However, not only have very few studies directly investigated interoceptive dimensions in samples of healthy adolescents and adults, but the existing evidence is also mixed (for an overview, see [13, 14]). For example, May et al. [19] reported different neural activity but no differences at the behavioural level between 16 adolescents (15-17 yrs), 19 young adults (20-28 yrs) and 19 mature adults (29-55 yrs) in an interoceptive task probing soft touch. Yang et al. [20], instead, found higher interoceptive accuracy in a sample of 50 adolescents (12-16 yrs) as compared to a sample of 50 adults (23-54 yrs), a finding that is in contrast with the idea that there is a disruption of interoception during adolescence [14]. However, considering that physiological ageing is associated with a reduction in interoception [13,14, 21] and that this last study included a sample of adults with a broad age range, it is difficult to determine whether adolescents truly performed better on the task, or whether the effect was due to the inclusion of middle-aged adults in the comparison group. In this study, a subsample of participants (35 adolescents and 21 adults) was also given a measure of ISe (i.e., Multidimensional Assessment of Interoceptive Awareness; [22]), and, in this case, adolescents showed a lower tendency to actively listen to the body for insight.

Qualitative findings also suggest that adolescents may experience body awareness in highly individualised ways, shaped by both bodily changes and psychological development. For instance, Pérez-Peña et al. [23] found that adolescents and young adults described their interoceptive experiences as fluctuating and context-dependent, often reflecting their struggles in interpreting bodily cues during times of emotional stress or social pressure. Such subjective accounts further emphasize the need to investigate interoceptive development through both quantitative and qualitative lenses. Thus, to advance our understanding of interoception development, the current study investigated multiple interoceptive dimensions during adolescence, analysing possible differences from the adult pattern of development. To these aims, healthy adolescents, whose ages ranged between 12 and 14 years, and adults, whose ages ranged between 25 and 34 years, performed a protocol that included measures of IAcc, ISe and IAw. Adults in this age range represent an optimal comparison group, avoiding confounding effects related to ageing processes. Indeed, several studies have suggested a regular decline in several cognitive skills (e.g., speed of processing, working memory, and long-term memory) starting from the 20s (see [24-26]), including interoceptive processing (for an overview see [21]).

## Materials and Methods

### Participants

Fifty-four typically developing adolescents (33 female participants, 21 male participants; mean age = 13.3 years, SD =0.75, range 12-14 years) and fifty adults (24 female participants, 26 male participants; mean age = 27.7 years, SD = 2.6, range 25-34 years) participated in this study.

A total of 50 adults and 54 adolescents were recruited based on sample sizes used in previous studies (e.g.,[20]). Additionally, a sensitivity power analysis was performed in G*Power 3.1.9.7 [27] for a two-sample t test, two-tailed, with α = .05 and desired power = .80. The analysis showed that the study was powered to detect effects as small as d = 0.56. Thus, any medium-to-large differences between adolescents and adults should have been detectable.

All participants were native Italians from an urban context in southern Italy. Adolescents were recruited from state schools in Calabria (Italy), while young adults were recruited by word of mouth.

All recruited participants showed normal reasoning ability according to the Italian norms of the Raven’s Colored Progressive Matrices (RCPM, [28,29]) or of Raven’s Standard Progressive Matrices (RSPM, for participants aged from 12 to 13 years; [30]) and had normal or corrected to normal vision and no history of neurological or psychiatric conditions.

Before taking part in the study, all adult participants provided written informed consent. Adolescents’ assent was received before the investigation, and their parents gave written informed consent. Participants were recruited between 9 April 2022 and 22 December 2023. The study was approved by the local ethics committee (Calabria Region Ethical Committee, Catanzaro, Italy) in accordance with the criteria laid down in the 1964 Declaration of Helsinki.

## Behavioral Testing

### Assessment of the Interoceptive Accuracy

A Heartbeat Monitoring Task (HMT) [31] was used to assess IAcc. Participants, placed in a comfortable position, were invited to relax, close their eyes, and focus on bodily sensations, and were told: “When you hear a voice say “*go*” start counting your heartbeats silently; when you hear “*stop*”, stop counting and tell me the exact number of heartbeats you counted”. They were also instructed not to move during the task and not to perform physical manipulations that could facilitate the detection of the pulse (for example, feeling the beat by testing the pulse). This task was repeated six times, using, for the adult group, time intervals of 25, 35 and 45 seconds separated by two standard rest periods of 20 seconds; shorter intervals of 15, 20 and 18 seconds were used for the adolescents [32].

To be sure that the instructions given to participants were clear, they were given a short training interval (10 seconds).

Meantime, real heart activity was recorded using a Bluetooth heart rate monitor (Polar Verity sense, Kempele, Finland), a mobile device that allows easy and non-invasive recording. The heartbeat signals of each participant were recorded and, through comparison with their count made by the participant, the IAcc was calculated. For each trial, an accuracy score was derived (using the formula of Garfinkel et al. [9]): 1 - (|n real beats - n counted beats|) / ((n real beats + n counted beats)/2). The accuracy scores obtained were calculated as the average of the six trials, producing an average value for each participant [33]. The inclusion of the reported values (n counted beats) within the denominator prevented an overestimation of the accuracy of performance in people who showed high variance, particularly when more heartbeats were reported than recorded [9].

At the end of the task, participants were asked, “*In which part of your body did you feel your heartbeat during the previous task*?”. Then, an image of a body map was presented (adapted from [34]), and participants were asked to indicate the relevant body areas by circling them. The image also includes a box above the head with the label “nowhere”. Nine body districts were identified: head, right ear, left ear, neck, chest, abdomen, right hand and wrist, left hand and wrist and legs; each body district was assigned 1 when the participant indicated that a specific part associated with the perception of the heartbeat. Zero was assigned to those body districts that were not selected by the participants.

### Assessment of the Interoceptive Awareness

IAw was assessed by the correlation between the measure of IAcc and the degree of confidence in one’s ability to estimate the number of heartbeats in the HMT, expressed by the participant at the end of each trial, on a scale from 0 to 10, where 0 indicated “no perception of heartbeat” and 10 indicated “full perception of heartbeat” (for such methodology see [9]).

### Assessment of the Interoceptive Sensibility

ISe was evaluated considering measures targeting both momentary, state-like beliefs (i.e., confidence ratings), and global, trait-like interoceptive beliefs (i.e., ISe questionnaires; see [35]).

Specifically, for what attains the state-like beliefs, at the end of each HMT trial, the participants rated their confidence in their perceived accuracy of response on a scale from 0 to 10, where 0 indicated “no perception of heartbeat” and 10 indicated “full perception of heartbeat”. The task included six trials, and a mean confidence score was computed for each participant by averaging the confidence ratings across the six trials. Participants also completed an ISe questionnaire. Specifically, adult participants completed the Self-Awareness Questionnaire (SAQ; [36]), while adolescents completed the SAQ-C, an adaptation of the SAQ for children and adolescents [37]. The SAQ and the SAQ-C are self-report questionnaires composed of 35 items developed specifically to evaluate the frequency of common body feelings. Both versions have been validated in Italian. Items are clustered into two domains, one related to visceral feelings (e.g., “I feel my heart beat in my ears”) and the other to somatosensory feelings (e.g., “I feel my palms sweaty”).

Participants were asked to read each item carefully and to evaluate how often they experienced the described sensation; responses were reported on a five-point Likert scale ranging from never to always (0 = never; 1 = sometimes; 2 = often; 3 = very often; 4 = always). The total score is given by the sum of the responses of all items, providing a score range of 0 to 140. Higher scores indicate higher levels of ISe.

### Statistical analyses

To verify the normality of data distribution for accuracy scores, we used the Shapiro-Wilk test. Given the non-normal distribution observed in experimental variables, such as the score of the heartbeat monitoring task, and considering that the questionnaire probing interoceptive sensibility (SAQ) used Likert-style response items, providing ordinal data, non-parametric statistical analyses were performed.

Specifically, comparisons between the two age groups (adolescents: 12 to 14 years old *vs*. adults: 25 to 34 years old) on IAcc, IAw and ISe scores were performed using the Mann-Whitney U test. Effect sizes for Mann–Whitney U tests were reported using the rank-biserial correlation coefficient r_rb_.

A Chi-squared test was applied to analyse which part of the body was most used during the IAcc task.

Finally, correlation analyses were conducted to explore the relationship between various interoceptive dimensions within the adolescent and young adult groups. Specifically, Spearman’s correlations were performed to examine the associations between IAcc (Heartbeat Monitoring Task), IAw and ISe (i.e., mean confidence in the HMT and SAQ total score) scores within each age group.

## Results

Descriptive statistics for IAcc, IAw and ISe measures are reported in Table 1. Concerning IAcc, the Mann–Whitney U tests revealed only a marginal difference between the group of adolescents and adults in counting their heartbeats (U = 1055, p = .055; r_rb_ = .22), with adults exhibiting numerically higher IAcc on average (see Table 1). Instaed, adolescents exhibited a statistically significantly lower metacognitive awareness of their interoceptive ability compared to adults (U= 981, p= .033; r_rb_ = .23). Concerning the ISe, the Mann-Whitney U test showed a significant effect of age group on both the SAQ (U= 843, p < .001; r_rb_ =0.38) and on the average confidence in the HMT (U= 911, p = .004; r_rb_ =0.33), with adolescents reporting significantly higher scores than adults.

**Table 1.** Descriptive statistics for the interoceptive measures in the groups of adolescents and adults.

| **Adolescent group** |  |  |  |  |
| --- | --- | --- | --- | --- |
|  | *Interoceptive*<br>*Accuracy* | *Interoceptive*<br>*Awareness* | *Interoceptive*<br>*Sensibility* |  |
|  | HMT | ACCURACY-CONFIDENCE<br>CORRELATION | SAQ | CONFIDENCE-HMT |
| Mean<br>(SD) | 0.36<br>(0.4) | 0.002<br>(0.5) | 44.8<br>(19) | 7.53<br>(1.25) |
| Min -Max | -0.95 – 0.91 | -1 – 0.96 | 12-94 | 4.8 – 10 |
| **Adult group** |  |  |  |  |
|  | *Interoceptive*<br>*Accuracy* | *Interoceptive*<br>*Awareness* | *Interoceptive*<br>*Sensibility* |  |
|  | HMT | ACCURACY-CONFIDENCE<br>CORRELATION | SAQ | CONFIDENCE-HMT |
| Mean<br>(SD) | 0.50<br>(0.4) | 0.24<br>(0.4) | 33.3<br>(14.3) | 6.57<br>(1.6) |
| Min- Max | -0.54 – 0.95 | -0.86 – 0.97 | 10-73 | 2.50– 9.33 |

Note: HMT, Heartbeat monitoring task; SAQ, Self-Awareness Questionnaire.

Correlation analyses showed no significant associations between the different interoceptive dimensions in both age groups (for *adolescents*, see Table 2; for *adults*, see Table 3).

**Table 2.** Spearman correlation coefficients between the interoceptive measures in the adolescent group.

| Adolescent group |  |  |  |  |  |
| --- | --- | --- | --- | --- | --- |
|  |  | *Interoceptive Accuracy* | *Interoceptive Awareness* | *Interoceptive Sensibility* |  |
|  |  | HMT | ACCURACY-CONFIDENCE CORRELATION | SAQ | CONFIDENCE-HMT |
| Interoceptive Accuracy | $r_{rho}$ | - | -0.07 | -0.06 | 0.18 |
| | $p$ | | .62 | .68 | .20 |
| Interoceptive Awareness | $r_{rho}$ | - | - | 0.16 | -0.04 |
| | $p$ | | | .27 | .81 |

Note: HMT, Heartbeat monitoring task; SAQ, Self-Awareness Questionnaire.

**Table 3.**
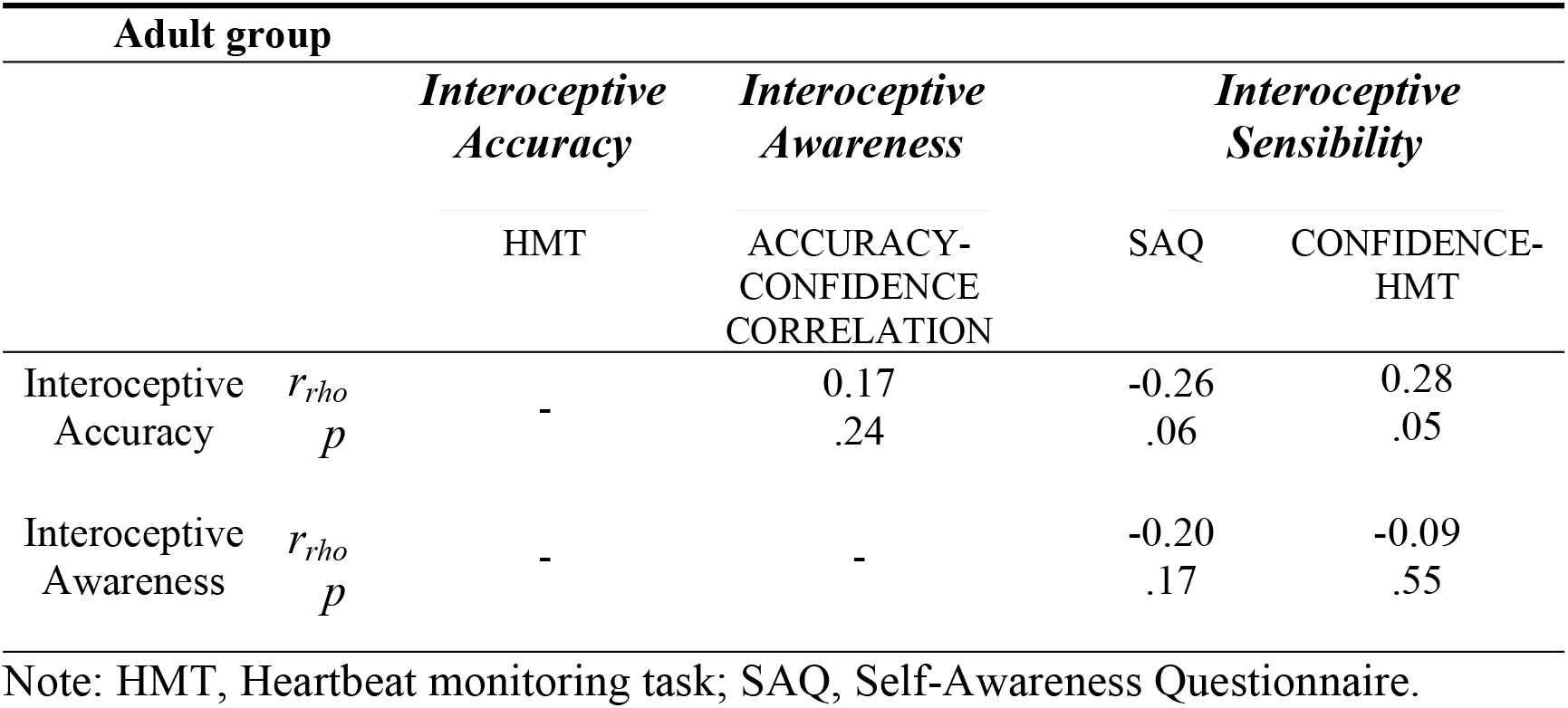
Spearman correlation coefficients between the interoceptive measures in the adult group.

Concerning the two ISe measures, we found no significant associations between the SAQ and the confidence rating in the HMT, both in adolescents (ISe-SAQ and Ise-confidance, r_rho_ = .05, p= .517) and in adults (ISe-SAQ and Ise-confidance, r_rho_ = .104, p= .471).

A chi-squared goodness of fit test was performed separately for adolescents and adults to examine which body parts were used most frequently during the IAcc task.

For adolescents, the distribution of selected body parts was significantly different from a uniform distribution (χ^2^(9) = 146, p < .001). The most commonly used body part was the chest (49.35%), followed by the right hand/wrist (14.29%) and the neck (10.39%). A similar pattern emerged for adults, χ^2^(9) = 88.0, p < .001, with the chest again being the most frequently selected body part (36.25%), followed by the right hand/wrist (17.5%) and left hand/wrist (16.25%).

To compare the distribution of body part selection between the two age groups (adolescent vs. adults), a Chi-squared test of independence was performed. Body-part selection did not differ significantly across age groups (χ^2^(8) = 8.63, p = .374). This suggests that the body parts used during the HMT was not significantly different between adolescents and adults.

## Discussion

The study examined interoceptive functioning across three distinct dimensions - accuracy, sensibility and awareness - in adolescents, with a focus on potential developmental differences when compared to adults.

While no significant group differences emerged in IAcc, there was a marginal trend suggesting slightly better performance in adults. More robust was the difference in IAw, with adults demonstrating significantly higher metacognitive insight into their bodily signals. On the other hand, adolescents scored higher on both measures of ISe: the SAQ and on confidence ratings during the heartbeat task. This pattern highlights a dissociation between the subjective experience of bodily awareness and the metacognitive ability to evaluate it.

The absence of a clear group difference in IAcc contrasts with previous findings by Yang et al. [20], who reported a higher level of accuracy in adolescents. This difference may be due to the broader age range of the adult sample by [20], which included individuals into middle adulthood, where interoceptive accuracy is thought to decline. By limiting our adult cohort to 25–34 years, we minimised this age-related confound and found that heartbeat-counting accuracy is comparable across late adolescence and early adulthood. Methodological differences (e.g., kind of task) may also have contributed to the discrepant results.

Regarding IAw, the finding of lower metacognitive insight in adolescents is in line with developmental models of metacognition (e.g., [38]) and supports the idea that IAw matures later than the basic ability to detect bodily signals. Conversely, the higher self-reported ISe in adolescents may reflect the heightened bodily attention characteristic of adolescence due to pubertal changes [13] or psychosocial factors. This finding is also interesting in light of studies that suggested that an exaggerated interoceptive sensibility can be dysfunctional (see [11]). Indeed, this increased tendency to notice internal bodily signals may be physiological, but can become fertile ground for the onset of mental health disorders (e.g., anxiety and depression).

The lack of significant correlations between interoceptive dimensions in both age groups replicates previous findings (for adolescents see [18]; for adults see [9]), and lends further support to the three-dimensional model of interoception proposed by [9]. Our data reinforce the idea that these dimensions are relatively independent and should not be interpreted as reflecting a unified construct.

In our data, even within each age group, objective performance did not correlate with subjective sensibility or metacognitive awareness, confirming that metacognitive or subjective insight into internal states does not necessarily align with actual IAcc. This underscores the need for a more nuanced approach when assessing interoceptive abilities.

Additionally, our findings revealed no correlation between the two ISe measures, that is, the SAQ scores and confidence ratings, within either group, in line with the idea that these measures probe different ISe aspects [35]. Indeed, while the SAQ captures a general, habitual focus on bodily signals, confidence ratings may reflect a momentary, context-dependent judgment of interoceptive certainty [35]. In developmental contexts, this is particularly relevant, as adolescents might report increased general interoceptive sensibility due to physical and emotional changes, without this necessarily translating into higher confidence in specific interoceptive tasks.

Regarding body localisation during the IAcc task, both adolescents and adults most frequently relied on the chest, followed by the wrists and neck. These patterns significantly deviated from a uniform distribution, indicating consistent preferences for certain bodily areas. However, the similarity between age groups in body part selection suggests that both adolescents and adults use comparable perceptual strategies when attending to internal sensations. This may reflect shared physiological or conceptual representations of interoceptive cues, such as the heartbeat, regardless of developmental stage.

Altogether, the findings contribute to a more comprehensive understanding of interoception in adolescence. They indicate that while basic detection of bodily signals may already be well established, the ability to reflect on or interpret these sensations continues to develop. The observed mismatch between heightened bodily focus (ISe) and lower metacognitive awareness (IAcc) in adolescents may have implications for emotional processing and vulnerability to psychological distress.

Given the role of interoception in emotion regulation and psychopathology, these results suggest that interventions aimed at adolescents could benefit from fostering the ability to accurately evaluate and understand internal states.

Despite its contributions, this study has some limitations. The cross-sectional design prevents us from inferring developmental trajectories, and our sample size, although adequate, could be increased to improve statistical power. The validity of the HMT has been questioned, as it may reflect participants’ estimation of their heart rate rather than their actual ability to feel the heartbeats [39 – 41]. Also, our IAcc and IAw measures exclusively targeted the cardiac modality.

Future studies should consider longitudinal designs and additional interoceptive measures that consider different organ systems, including the cardiac, gastric, and respiratory systems (for an overview, see [42]), to enhance our understanding of interoceptive development.

## References

1. Craig AD. How do you feel? Interoception: the sense of the physiological condition of the body. Nat Rev Neurosci. 2002;3(8):655–66. doi:10.1038/nrn894

2. Garfinkel SN, Critchley HD. Interoception, emotion and brain: new insights link internal physiology to social behaviour. Commentary on: “Anterior insular cortex mediates bodily sensibility and social anxiety” by Terasawa et al. (2012). Soc Cogn Affect Neurosci. 2013;8(3):231–4. doi:10.1093/scan/nss140

3. Canino S, Torchia V, Gaita M, Raimo S, Palermo L. Linking the inner and outer mental representations of the body to social cognition skills: A systematic review and meta-analysis. Neuropsychologia. 2024;204:108989. doi:10.1016/j.neuropsychologia.2024.108989

4. Critchley HD, Garfinkel SN. Interoception and emotion. Curr Opin Psychol. 2017;17:7–14. doi:10.1016/j.copsyc.2017.04.020

5. Füstös J, Gramann K, Herbert BM, Pollatos O. On the embodiment of emotion regulation: interoceptive awareness facilitates reappraisal. Soc Cogn Affect Neurosci. 2013;8(8):911–7. doi:10.1093/scan/nss089

6. Khalsa SS, Adolphs R, Cameron OG, Critchley HD, Davenport PW, Feinstein JS, et al. Interoception and mental health: a roadmap. Biol Psychiatry Cogn Neurosci Neuroimaging. 2018;3(6):501–13. doi:10.1016/j.bpsc.2017.12.004

7. Raimo S, Martini M, Guariglia C, Santangelo G, Trojano L, Palermo L. Editorial: Community series in body representation and interoceptive awareness: cognitive, affective, and social implications. Front Psychol. 2023a;14:1256811. doi:10.3389/fpsyg.2023.1256811

8. Tsakiris M, Critchley H. Interoception beyond homeostasis: affect, cognition and mental health. Philos Trans R Soc Lond B Biol Sci. 2016;371(1708):20160002. doi:10.1098/rstb.2016.0002

9. Garfinkel SN, Seth AK, Barrett AB, Suzuki K, Critchley HD. Knowing your own heart: distinguishing interoceptive accuracy from interoceptive awareness. Biol Psychol. 2015;104:65–74. doi:10.1016/j.biopsycho.2014.11.004

10. Baiano C, Job X, Santangelo G, Auvray M, Kirsch LP. Interactions between interoception and perspective-taking: current state of research and future directions. Neurosci Biobehav Rev. 2021;130:252–62. doi:10.1016/j.neubiorev.2021.08.007

11. Garfinkel SN, Tiley C, O’Keeffe S, Harrison NA, Seth AK, Critchley HD. Discrepancies between dimensions of interoception in autism: implications for emotion and anxiety. Biol Psychol. 2016;114:117–26. doi:10.1016/j.biopsycho.2015.12.003

12. Scarpazza C, Zangrossi A, Huang YC, Sartori G, Massaro S. Disentangling interoceptive abilities in alexithymia. Psychol Res. 2022;86(3):844–57. doi:10.1007/s00426-021-01538-x

13. Murphy J, Brewer R, Catmur C, Bird G. Interoception and psychopathology: a developmental neuroscience perspective. Dev Cogn Neurosci. 2017;23:45–56. doi:10.1016/j.dcn.2016.12.006

14. Carr L, Donaghy R, Brewer R. Interoception across the lifespan.In: Murphy J, Brewer R, editors. Interoception. Cham: Springer; 2024. doi:10.1007/978-3-031-68521-7_10

15. Zvolensky MJ, Smits JAJ, editors. Anxiety in health behaviors and physical illness. Springer; 2007.

16. Li D, Zucker NL, Kragel PA, Covington VE, LaBar KS. Adolescent development of insula-dependent interoceptive regulation. Dev Sci. 2017;20(5):e12438. doi:10.1111/desc.12438

17. Price CJ, Hooven C. Interoceptive awareness skills for emotion regulation: theory and approach of mindful awareness in body-oriented therapy (MABT). Front Psychol. 2018;9:798. doi:10.3389/fpsyg.2018.00798

18. Mai S, Wong CK, Georgiou E, Pollatos O. Interoception is associated with heartbeat-evoked brain potentials (HEPs) in adolescents. Biol Psychol. 2018;137:24– 33. doi:10.1016/j.biopsycho.2018.06.007

19. May AC, Stewart JL, Tapert SF, Paulus MP. The effect of age on neural processing of pleasant soft touch stimuli. Front Behav Neurosci. 2014;8:52. doi:10.3389/fnbeh.2014.00052

20. Yang HX, Zhou HY, Wei Z, Wan GB, Wang Y, Wang YY, et al. Multidimensional interoception and autistic traits across life stages: evidence from a novel eye-tracking task. J Autism Dev Disord. 2022;52:2644–55. doi:10.1007/s10803-021-05155-w

21. Pfeifer G, Cawkwell S. Interoceptive ageing and the impact on psychophysiological processes: a systematic review. Int J Psychophysiol. 2025;207:112483. doi:10.1016/j.ijpsycho.2024.112483

22. Mehling WE, Price C, Daubenmier JJ, Acree M, Bartmess E, Stewart A. The multidimensional assessment of interoceptive awareness (MAIA). PLoS One. 2012;7(11):e48230. doi:10.1371/journal.pone.0048230

23. Pérez-Peña M, Notermans J, Petit J, Van der Gucht K, Philippot P. Body aware: adolescents’ and young adults’ lived experiences of body awareness. Psychol Belg. 2024 Aug 12;64(1):108–28. doi:10.5334/pb.1295

24. Salthouse TA. The processing-speed theory of adult age differences in cognition. Psychol Rev. 1996;103(3):403. doi:10.1037/0033-295X.103.3.403

25. Park DC, Reuter-Lorenz P. The adaptive brain: aging and neurocognitive scaffolding. Annu Rev Psychol. 2009;60:173–96. doi:10.1146/annurev.psych.59.103006.093656

26. Park DC, Lautenschlager G, Hedden T, Davidson NS, Smith AD, Smith PK. Models of visuospatial and verbal memory across the adult life span. Psychol Aging. 2002;17(2):299–320. doi:10.1037/0882-7974.17.2.299

27. Faul F, Erdfelder E, Buchner A, Lang AG. Statistical power analyses using G*Power 3.1: tests for correlation and regression analyses. Behav Res Methods. 2009;41:1149–60.

28. Basso A, Capitani E, Laiacona M. Raven’s coloured progressive matrices: normative values on 305 adult normal controls. Funct Neurol. 1987;2(2):189–94.

29. Belacchi C, Scalisi TG, Cannoni E, Cornoldi C. Matrici progressive di Raven forma colore (CPM-47): manuale d’uso e standardizzazione italiana. Firenze: Giunti Organizzazioni Speciali; 2008.

30. Picone L, Orsini A, Pezzuti L. Raven’s Standard Progressive Matrices: contribution to Italian standardization for subjects between ages 6 and 18. BPA Appl Psychol Bull. 2017;65(280).

31. Schandry R. Heart beat perception and emotional experience. J Psychophysiol. 1981;18:483–8. doi:10.1111/j.1469-8986.1981.tb02486.x

32. Koch A, Pollatos O. Cardiac sensitivity in children: sex differences and its relationship to parameters of emotional processing. J Psychophysiol. 2014;51(9):932– 41. doi:10.1111/psyp.12233

33. Hart N, McGowan J, Minati L, Critchley HD. Emotional regulation and bodily sensation: interoceptive awareness is intact in borderline personality disorder. J Pers Disord. 2013;27(4):506–18. doi:10.1521/pedi_2012_26_049

34. Plans D, Ponzo S, Morelli D, Cairo M, Ring C, Keating CT, et al. Measuring interoception: the phase adjustment task. Biol Psychol. 2021;165:108171. doi:10.1016/j.biopsycho.2021.108171

35. Suksasilp C, Garfinkel SN. Towards a comprehensive assessment of interoception in a multi-dimensional framework. Biol Psychol. 2022;168:108262. doi:10.1016/j.biopsycho.2022.108262

36. Longarzo M, D’Olimpio F, Chiavazzo A, Santangelo G, Trojano L, Grossi D. The relationships between interoception and alexithymic trait. The Self-Awareness Questionnaire in healthy subjects. Front Psychol. 2015;6:1149. doi:10.3389/fpsyg.2015.01149

37. Raimo S, Iona T, Di Vita A, Boccia M, Torchia V, Canino S, et al. The interoceptive sensibility in middle childhood: the Italian validation of the Self-Awareness Questionnaire. Eur J Dev Psychol. 2023b;21(1):138–53. doi:10.1080/17405629.2023.2250121

38. Weil LG, Fleming SM, Dumontheil I, Kilford EJ, Weil RS, Rees G, et al. The development of metacognitive ability in adolescence. Conscious Cogn. 2013;22(1):264–71. doi:10.1016/j.concog.2013.01.004

39. Addabbo M, Milani L. Measuring interoception from infancy to childhood: a scoping review. Neurosci Biobehav Rev. 2025 Jun;173:106161. doi:10.1016/j.neubiorev.2025.106161. PMID:40245971.

40. Corneille O, Desmedt O, Zamariola G, Luminet O, Maurage P. A heartfelt response to Zimprich et al. (2020), and Ainley et al. (2020)’s commentaries: acknowledging issues with the HCT would benefit interoception research. Biol Psychol. 2020;152:107869. doi:10.1016/j.biopsycho.2020.107869

41. Desmedt O, Luminet O, Corneille O. The heartbeat counting task largely involves non-interoceptive processes: evidence from both the original and an adapted counting task. Biol Psychol. 2018;138:185–8. doi:10.1016/j.biopsycho.2018.09.004

42. Desmedt O, Luminet O, Walentynowicz M, Corneille O. The new measures of interoceptive accuracy: a systematic review and assessment. Neurosci Biobehav Rev. 2023;153:105388.

